# Parotid glands have a dysregulated immune response following radiation therapy

**DOI:** 10.1101/2023.11.27.568872

**Authors:** Jordan A. Gunning, Kristy E. Gilman, Tiffany M. Zúñiga, Richard J. Simpson, Kirsten H. Limesand

## Abstract

Head and neck cancer treatment often consists of surgical resection of the tumor followed by ionizing radiation (IR), which can damage surrounding tissues and cause adverse side effects. The underlying mechanisms of radiation-induced salivary gland dysfunction are not fully understood, and treatment options are scarce and ineffective. The wound healing process is a necessary response to tissue injury, and broadly consists of inflammatory, proliferative, and redifferentiation phases with immune cells playing key roles in all three phases. In this study, select immune cells were phenotyped and quantified, and certain cytokine and chemokine concentrations were measured in mouse parotid glands after IR. Further, we used a model where glandular function is restored to assess the immune phenotype in a regenerative response. These data suggest that irradiated parotid tissue does not progress through a typical inflammatory response observed in wounds that heal. Specifically, total immune cells (CD45+) decrease at days 2 and 5 following IR, macrophages (F4/80+CD11b+) decrease at day 2 and 5 and increase at day 30, while neutrophils (Ly6G+CD11b+) significantly increase at day 30 following IR. Additionally, radiation treatment reduces CD3-cells at all time points, significantly increases CD3+/CD4+CD8+ double positive cells, and significantly reduces CD3+/CD4-CD8-double negative cells at day 30 after IR. Previous data indicate that post-IR treatment with IGF-1 restores salivary gland function at day 30, and IGF-1 injections attenuate the increase in macrophages, neutrophils, and CD4+CD8+ T cells observed at day 30 following IR. Taken together, these data indicate that parotid salivary tissue exhibits a dysregulated immune response following radiation treatment which may contribute to chronic loss of function phenotype in head and neck cancer survivors.

## Introduction

Over 600,000 new cases of head and neck cancer are diagnosed annually, with treatment consisting of surgical resection of the tumor, followed by radiation with or without chemotherapy(1, 2). Radiation treatment for head and neck cancer often exposes nondiseased tissues to ionizing radiation and can lead to deleterious side effects. Salivary gland hypofunction and xerostomia are common and often chronic side effects of radiotherapy in head and neck cancer patients and can result in increased dental caries and mucositis, as well as difficulty eating, speaking, and swallowing(3). Currently available treatments have severe side effects frequently resulting in cessation of treatment, or only temporarily relieve the symptoms and do not provide permanent restoration of salivary gland function, as defined by an improvement in salivary flow rate. Gaining a better understanding of the mechanisms involved could lead to more efficacious treatments and improvements in quality of life for head and neck cancer survivors.

Wound healing is a highly coordinated physiological reaction to tissue injury and involves complex interactions between multiple cell types, mediators and signaling molecules, and the surrounding vasculature(4–7). The immune system plays a pivotal role in orchestrating wound healing, and infiltration of immune cells and production of inflammatory mediators in damaged tissue is essential for tissue regeneration and restoration of function. Immune infiltration and an increase in cytokines are observed in several tissues following IR, including lung, intestine, and the heart(8–11). Neutrophils are granular innate immune cells recruited to the site of injury within hours in response to damage-associated molecular patterns (DAMPs), including extracellular ATP, reactive oxygen species, and heat-shock proteins(12–16). Macrophages are highly plastic cells of the innate immune system with roles in phagocytosing dead cells, including neutrophils, and bacteria in the injured tissue, secreting cytokines, and inducing T cell activation(17–19). Macrophages are essential for all stages of wound healing as well as full restoration of tissue function, and various methods of *in vivo* macrophage depletion show significant impairments in wound healing(20, 21). Chemokines and chemoattractants are released from resident immune cells at the site of injury and stimulate a gradient to induce leukocyte migration, which is necessary for leukocyte extravasation to occur. Specific pro- and anti-inflammatory cytokines, including IL-6 and IL-10, have been shown to be necessary for proper wound healing, and suppression of these with monoclonal antibodies or genetic disruption inhibits would healing(22–24). Lymphocytes are the last immune cell type to infiltrate wounded tissue and remain for at least 30 days following an injury; however, lymphocytes are highly radiosensitive and radiotherapy often causes immunosuppression and rapid clearing of lymphocytes from the radiation site(25–27). The role of lymphocytes in healthy tissue regeneration is not well understood, but it is thought that CD4+ T cells are the most important lymphocytic cell type in wound healing(28, 29). Certain lymphocytes, such as FoxP3+ regulatory T cells, have immunosuppressive properties in wound healing that dampen the inflammatory response and help tissue heal fully(30, 31). It is well documented that acute inflammation and immune activity are essential to proper healing following injury, but persistent inflammation inhibits progression through the wound healing cascade(32).

Interestingly, treating mice with indomethacin, a cyclooxygenase (COX) inhibitor, reduces pro-inflammatory prostaglandin concentrations in parotid tissue and restores salivary gland function at day 30 when given at days 3, 5, and 7 post-IR(33). However, no improvement in glandular function is seen when indomethacin is administered prior to IR(33). This suggests that an acute inflammatory response within the first two days following IR may be necessary for parotid salivary glands to recover (33). Moreover, treatment with insulin-like growth factor-1 (IGF-1) restores salivary flow rates when given at days 4-7 post-IR by attenuating compensatory proliferation, maintaining activation of aPKCζ, and increasing expression and secretion of amylase; however, the effect of IGF-1 treatment on the immune response after IR is not known(2, 34–41). Macrophage infiltration is observed in multiple tissues following IR, and is often correlated with fibrosis, particularly in lung and cardiac tissue(9–11, 42). Resident macrophages in the submandibular gland have been found to decrease significantly following IR, and mRNA markers of macrophages are significantly decreased in the swine parotid gland at 1 and 5 weeks post IR(43, 44). Interestingly, these studies also reported that treatments used to restore salivary flow also restore macrophage populations, suggesting they may play an important role in salivary gland regeneration. Additionally, concentrations of pro-inflammatory cytokines as well as pro-inflammatory gene expression have been shown to increase following IR; TNF-α and IL-1β increase in splenocytes and intestines, and TNF-α increases in the lung(45–47). Some studies report treatment with bioactive compounds that inhibit cytokine production or direct inhibition of cytokines is radioprotective by attenuating radiation-induced apoptosis(47, 48). In the submandibular gland, IL-6 gene expression significantly increased at 3 hours and 14 days following 13 Gy IR(49), and another study found increased IL-6 and TNF-α mRNA expression and protein levels at week 5 after IR in swine parotid glands(50). Furthermore, treatment with exogenous IL-6 as well as genetic IL-6 ablation restored salivary gland function, as measured by improved saliva flow, at 8 weeks after IR(49). These data leave the inflammatory response to radiation in salivary glands unclear; thus, improving the understanding of the immune and inflammatory response following IR presents an opportunity to possibly identify therapeutic targets that reverse the loss of function.

In the present study, certain phagocytic and lymphocytic immune cells in the parotid gland were phenotyped and quantified at acute and chronic time points following IR compared to non-irradiated control mice. Additionally, cytokine and chemokine concentrations were determined in the parotid gland following IR. Given the well-established role of the immune system and acute inflammation in a normal wound healing response, we examined the possibility that irradiated parotid glands would exhibit an abnormal immune phenotype as compared to untreated mice due to their dysregulated proliferative response and lack of regeneration.

## Methods

### Mice

All mice were maintained in strict accordance with Institutional Animal Care and Use Committee (IACUC) regulations at the University of Arizona with protocols approved by IACUC. FVB/NJ mice were purchased from Jackson Labs (Bar Harbor, ME). Mice were housed in vented cages with *ad libitum* access to food and water following 12-hour light/dark cycles. Mice were sedated with intraperitoneal injection of a ketamine/xylazine mixture (70 mg/kg, 10 mg/mL) prior to radiation and salivary gland dissections. Mice were constrained in 50 mL conical tubes with breathing holes for radiation treatment. The head and neck region was exposed to a single 5 Gy dose of X-ray irradiation (RadSource, Kansas City, MO) while the rest of the body was shielded with >6mm of lead. For all experiments, parotid glands were rigorously cleared of lymph nodes and in the rare instance that lymph node contamination was detected by flow cytometry, these samples were not included in the analysis.

### Insulin-like growth factor (IGF)-1 injections

Mice were given a maximum of three IGF-1 doses (5μg/mouse) via tail-vein injection 24-hours apart on days 4 to 6 following radiation treatment. For analysis at day 5, mice received only one injection at day 4 post IR and salivary glands were harvested 24 hours later. For analysis at day 30, mice received three IGF-1 injections at days 4, 5, and 6 after IR and parotid glands were dissected on day 30 following radiation treatment. For IGF-1 analysis without radiation treatment, mice received three IGF-1 injections and parotid glands were removed 24 hours or 24 days after the last injection to mimic a day 30 timepoint.

### Labeling with monoclonal antibodies against surface antigens

An eight-color direct immunofluorescence protocol was used to label digested murine salivary glands with the following monoclonal antibodies: VioBlue-labeled Ly-6G, VioGreen-labeled CD11b, APC-labeled F4/80, APC-Vio 770-labeled CD45, VioBlue-labeled CD4, VioGreen-labeled CD8a, FITC-labeled CD3, and PE-Vio 770-labeled CD49b (Miltenyi Biotec, Auburn, CA) and analyzed by flow cytometry (Miltenyi Biotec MacsQuant, Auburn, CA). In brief, a single cell suspension was prepared by homogenizing tissue in media supplemented with 1 mg/mL collagenase and hyaluronidase, followed by an incubation at 37°C for 20 minutes, washing with EGTA, and nutating in EGTA for 10 minutes. Samples were then passed through a 70 µM filter screen to break up any remaining clumps, and cells were washed twice with 1X PBS before being passed through a 20 µM filter. Aliquots of 500 µL of a single cell suspension of salivary tissue were incubated with 10 µL of FcR mouse blocking reagent for 10 minutes (Miltenyi Biotec), followed by 0.5 or 1 µL of each mAb for 30 minutes at room temperature. The samples were then incubated with 2 mL of diluted RBC lysis buffer (Miltenyi Biotec, Auburn, CA) for 15 minutes at room temperature, washed with PBS, and resuspended in 200 µL PBS prior to flow cytometry analysis. The samples were run with propidium iodide (PI), which stains for dead cells, to determine cell viability. Data for a majority of figures was collected on a Miltenyi Biotec MACSQuant. The data shown in supplemental figure 6 was collected on a BD FACSCanto II, due to unavailability of the original cytometer. Following data acquisition, files were analyzed manually using FlowLogic software and percentages of relevant cell types were tabulated for statistical analysis. Single color controls performed with anti-REA comp beads (Miltenyi Biotec, Auburn, CA) were used to guide gating strategy, and single color control flow cytometry plots are available in supplemental figures 1 and 3. Our gating strategy is outlined in supplemental figures 2, 4, 7, and 8.

### Immunofluorescent staining

Salivary gland tissue was removed and immediately fixed in 10% formalin (ThermoFisher Scientific, Waltham, MA) for 24 hours. Tissues were shipped to IDEXX Laboratories for embedding and sectioning. Slides were baked for 20 minutes at 37°C, then rehydrated in Histoclear, graded ethanol (100-50%) and distilled water by performing 2 sequential washes in each solution for 5 minutes each. A permeabilization step in 0.2% Triton X-100 (MP Biomedicals, Santa Ana, CA) and 0.05% Tween20 (ThermoFisher Scientific, Waltham, MA) in 1X PBS was performed for 15 minutes prior to antigen retrieval, which was done by boiling samples in microwave for 15 minutes in 1 mM citric acid buffer (pH 6.0). The slides were left in citric acid buffer citric acid for 20 minutes at room temperature to cool. Slides were washed in PBS for 15 minutes and the tissues were outlined using a wax pen. Non-specific binding was blocked with 300 µL of 5% goat serum (EMD Millipore, Burlington MA) for 1 hour and then incubated with a 1:250 dilution of monoclonal rabbit F4/80 or 1:150 of rabbit Ly-6G (Cell Signaling, Danvers, MA) overnight at 4°C. Tissues were washed with 0.1% Triton X-100 in PBS, and then in PBS, and incubated in goat anti-rabbit AF 594-conjugated secondary antibody (1:500) (ThermoFisher Scientific, Waltham, MA) at room temperature for 1 hour. The slides were counterstained with DAPI (1 µg/mL) and mounted with ProLong gold antifade reagent (ThermoFisher Scientific, Waltham, MA). Fluorescent images were visualized on a Leica DM5500 Microscope System and digitally captured with a Pursuit 4 Megapixel CCD camera using Image Pro 7.0 software and morphometric analysis was performed with ImagePro 7.0 (Media Cybernetics, Rockville, MD).

### Cytokine array

Mice were untreated or exposed to 5Gy of radiation and parotid glands were excised at indicated timepoints (days 2, 7, 14, or 30) post-IR. Tissues were homogenized in PBS with protease inhibitor cocktail (30 µL/mL), PMSF (1mM) and Triton-X 100 (1%). Protein was quantified with the Pierce Coomassie Plus Bradford Assay (ThermoFisher Scientific, no. 23236, Waltham, MA) and 400 µg of tissue lysates were used per sample. Cytokine levels were determined with the Mouse Proteome Profiler XL Cytokine Array (R&D systems, no. ARY028, Minneapolis, MN) following the manufactures’ instructions with a 20-minute exposure to autoradiography film (Genesee, no. 30-810, Rochester, NY). Images were quantified using ImageJ software (NIH).

### Statistical analysis

Statistical analysis was done using GraphPad Prism 6 software (GraphPad Software, La Jolla, CA). To determine statistical significance between groups, a one-way ANOVA with Tukey’s post-hoc test was performed. Data is presented as samples ± SEM, and groups with different letter designations are significantly different from each other. A p value of <0.05 was considered to be significant.

## RESULTS

### Radiation leads to significant reductions in total immune cells and total macrophages in the parotid gland at days 2 and 5 following IR and an increase in macrophages and neutrophils at day 30 after IR

We first aimed to assess how radiation treatment influences the presence of phagocytic immune cells in murine salivary glands. Mice were treated with 5Gy IR and parotid tissue was dissected at days 2, 5, 14, and 30 following IR. Flow cytometry was performed on whole parotid tissue to quantify and phenotype total immune cells. Total viable immune cells (CD45+PI-) and viable macrophages (F4/80+CD11b+) were significantly decreased at days 2 and 5 post IR but return to untreated levels at days 14 post IR (Fig 1A, Fig 1B, Supplemental fig 2). Total viable immune cells remain unchanged from UT at day 30; however, macrophages are significantly elevated at day 30 following IR (Fig 1A, Fig 1B). Further, we observed that neutrophils (Ly6G+CD11b+) did not change at acute time points but were significantly increased at day 30 following IR (Fig 1C, Supplemental fig 2). We confirmed these data with immunofluorescent staining of parotid gland sections with anti-F4/80 and anti-Ly-6G and analysis by manual counting (Fig 1D-G). These data suggest radiation is causing a significant reduction in total immune cells and total macrophages in parotid glands at acute time points and thus, parotid glands appear to follow an atypical healing timeline.

**Fig 1.**
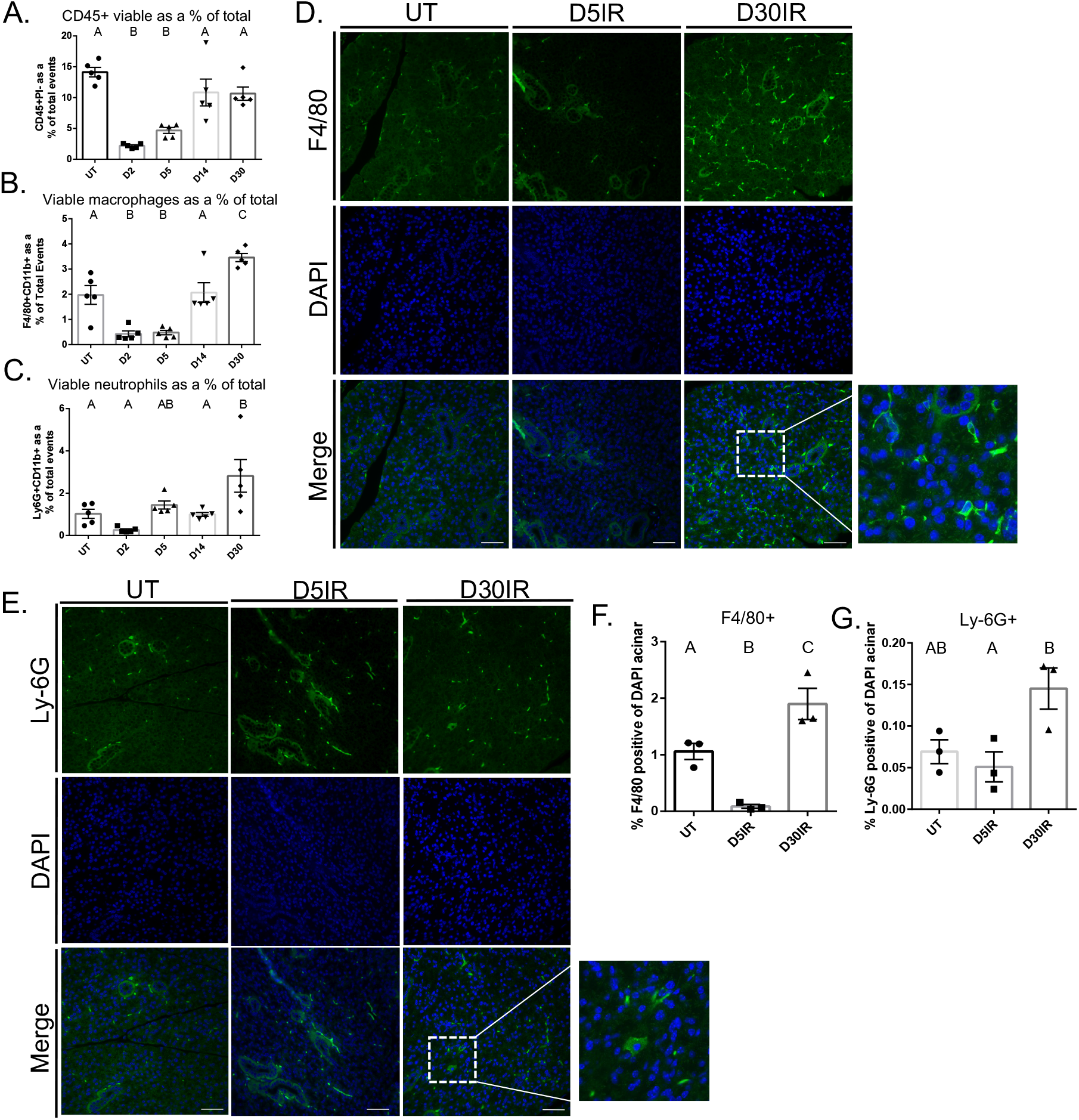
Total viable immune cells and macrophages decrease acutely while neutrophils and macrophage increase at day 30 in parotid glands after 5Gy IR. 4-6 week old FVB mice were treated with a single 5Gy dose of targeted head and neck irradiation. The parotid glands were dissected at days 2, 5, 14, and 30 following IR, prepared as a single cell suspension, and labeled with antibodies for phenotyping via flow cytometry. A) Viable total immune cells, defined as CD45+PI-. B) Viable F4/80+CD11b+ macrophages. C) Viable Ly-6G+CD11b+ neutrophils. D) Immunofluorescent staining with anti-F4/80 on FFPE salivary gland tissue. E) Immunofluorescent staining with anti-Ly-6G on FFPE salivary gland tissue. F) Quantification of F4/80 immunofluorescent staining determined by manual counting. G) Quantification of Ly-6G immunofluorescent staining determined by manual counting. Scale bars represent 50 µM. Data analyzed by one-way ANOVA with Tukey’s post-hoc test and represented as mean ± SEM. Treatment groups with the same letter are not statistically different from each other.

**Supplemental Fig 1. Single color controls for F4/80, Ly-6G, CD11b, and CD45.** Flow cytometry data were analyzed manually via FlowLogic. Gating strategy was guided by single color controls for A) CD45, B) CD11b, C) F4/80, D) Ly-6G using anti-REA comp beads.

**Supplemental Fig 2. Flow cytometry plots and gating strategy for phagocytic immune cells from IR damaged parotid glands.** Flow cytometry data was analyzed manually via FlowLogic. Viable CD45+PI-gated from total events, F4/80+CD11b+ macrophages gated from CD45+PI-, Ly-6G+CD11b+ gated from CD45+PI-in: A-C) UT mice, D-F) D2IR mice, G-I) D5IR mice, J-L) D14IR mice, and M-O) D30IR mice.

### PI+ total immune cells decrease at days 2, 5, and 14 post IR, and PI+ neutrophils significantly increase at day 30 post IR

Radiation induces a robust cell death response, and it is unclear if populations of viable immune cells decrease significantly due to cell death following radiation. Therefore, we sought to evaluate whether IR induces an increase in propidium iodide (PI)+ cells as a measure of immune cells undergoing cell death in the salivary gland and if this is concomitant with the decrease in viable immune cell populations. Total PI+ immune cells (CD45+PI+) were significantly decreased from untreated mice at days 2, 5, and 14 following IR, but were not significantly different at day 30 after IR (Fig 2A). PI+ macrophages were unchanged when compared to untreated at any time point measured (Fig 2B). Interestingly, PI+ neutrophils were unchanged at days 2, 5, and 14, and significantly increased at day 30 post IR (Fig 2D). Taken together, these data suggest that cell death is not responsible for the significant decreases in viable immune cell populations in parotid glands following IR.

**Fig 2.**
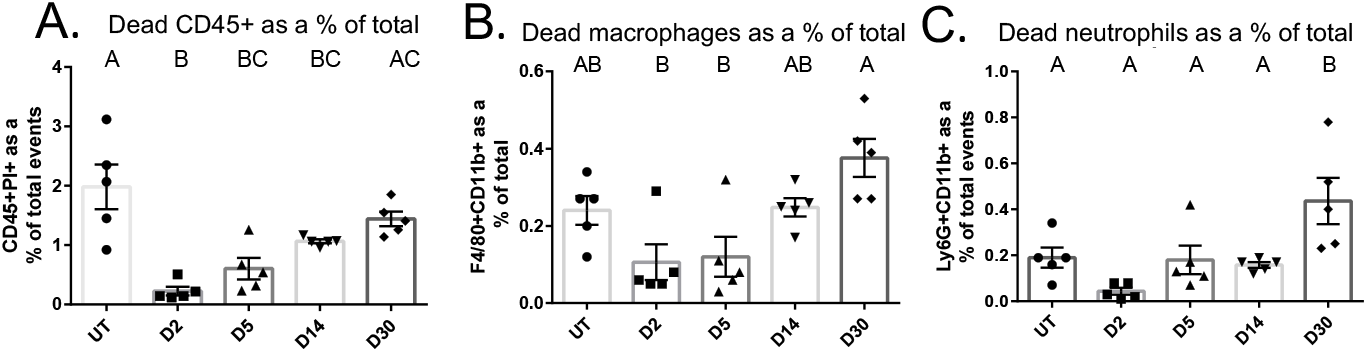
PI+ total immune cells are decreased at days 2, 5, and 14 following IR, and PI+ neutrophils are increased at day 30. Parotid glands were prepared as described in figure 1. A) Total PI+ immune cells, defined as CD45+PI+. B) PI+ F4/80+CD11b+ macrophages. C) PI+ Ly-6G+CD11b+ neutrophils. Data analyzed by one-way ANOVA with Tukey’s post-hoc test and represented as mean ± SEM. Treatment groups with the same letter are not statistically different from each other.

### Radiation significantly reduces CD3-cells and increases CD4+CD8+ double positive T cells at day 30 after IR

Next, we quantified and phenotyped lymphocytes in the parotid salivary gland following radiation treatment in order to characterize non-myeloid immune cells. Viable CD3+ T cells were not significantly different from untreated at any time point (Fig 3A, Supplemental fig 4). Further, viable CD3-cells were decreased at all time points following IR (Fig 3B, Supplemental fig 4). NK cells were not significantly different from untreated at any time point measured (Fig 3C, Supplemental fig 4). Both CD4+ and CD8+ single positive cells, gated from viable CD3+ cells, did not change following IR (Fig 3D, 3E, Supplemental fig 4). Remarkably, CD4+CD8+ double positive cells analyzed as a percentage of viable CD3+ T cells were significantly increased at day 30 following IR, while CD4-CD8-as a percentage of CD3+ T cells were decreased at day 30 (Fig 3F, 3G). These data further suggest that salivary glands have a dysregulated healing response as lymphocytes are vital for healing and typically increased at the site of injury through day 30(25).

**Fig 3.**
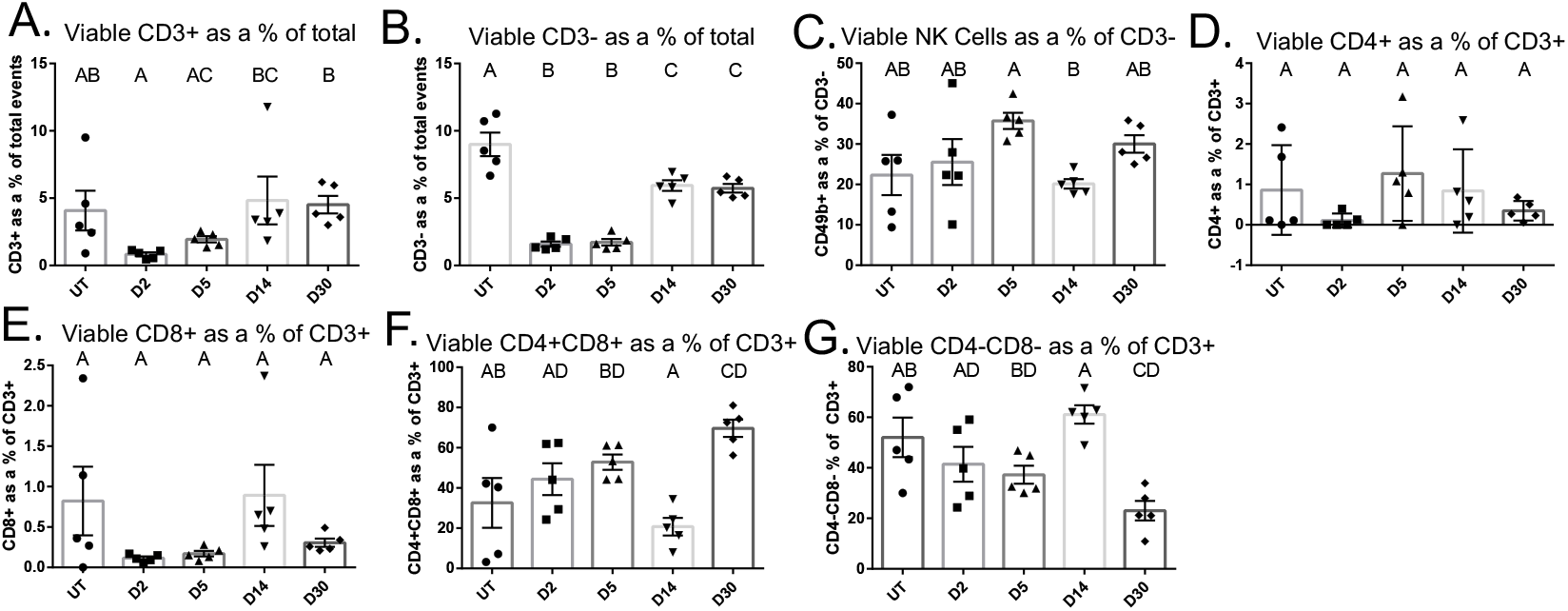
CD4+CD8+ double positive T cells significantly increase at day 30 following IR. Parotid glands were prepared as described in figure 1. A) Viable CD3+ T cells. B) Viable CD3-cells. C) Viable CD49b+ NK cells. D) Viable CD4+ T cells. E) Viable CD8+ T cells. F) Viable CD4+CD8+ double positive T cells. G) Viable CD4-CD8-double negative T cells. Data analyzed by one-way ANOVA with Tukey’s post-hoc test and represented as mean ± SEM. Treatment groups with the same letter are not statistically different from each other.

**Supplemental Fig 3. Single color controls for CD3, CD4, CD8, and CD49b antibodies.** Flow cytometry data were analyzed manually via FlowLogic. Gates were determined using isotype or single color controls for A) CD3, B) CD4, C) CD8, and D) CD49b.

**Supplemental Fig 4. Flow cytometry plots and gating strategy for lymphocytes in IR damaged parotid glands.** Flow cytometry data was analyzed manually via FlowLogic. CD3+ and CD3-gated from CD45+PI-, NK cells gated from CD3-cells, CD4+, CD8+, CD4-CD8-, CD4+CD8+ gated from CD3+ in: A-C) UT mice, D-F) D2IR mice, G-I) D5IR mice, J-L) D14IR mice, and M-O) D30IR mice.

### PI+ CD3+ T cells decrease at day 14 after IR while PI+ CD3-cells significantly decrease at all time points

Next, we analyzed PI+ lymphocytes in the parotid gland following IR. PI+ CD3+ T cells were significantly decreased at day 14 post IR, and PI+ CD3-cells were significantly decreased at all time points measured following radiation (Supplemental fig 5A, 5B). PI+ NK cells were significantly decreased at days 14 and 30 following IR (Supplemental fig 5C). PI+ CD4+, CD8+, CD4+CD8+, and CD4-CD8-T cells remain unchanged at all time points following radiation (Supplemental fig 5D-G). Similar to what we observed with phagocytic immune cells, these data suggest cell death is not responsible for the decreases in lymphocytic immune cells.

**Supplemental fig 5: PI+ CD3+ T cells decrease at day 14 after IR while PI+ CD3-cells decrease at all time points.** Parotid glands were prepared as described in figure 1. A) PI+ CD3+ T cells. B) PI+ CD3-cells. C) PI+ CD49b+ NK cells. D) PI+ CD4+ T cells. E) PI+ CD8+ T cells. F) PI+ CD4+CD8+ double positive T cells. G) PI+ CD4-CD8-double negative T cells. Data analyzed by one-way ANOVA with Tukey’s post-hoc test and represented as mean ± SEM. Treatment groups with the same letter are not statistically different from each other.

### IGF-1 administration significantly reduces IR-induced increases in viable macrophages and neutrophils 30 days post treatment

Subsequently, we sought to determine whether the inflammatory response is modulated in a model of glandular restoration post-IR. Previous studies have demonstrated that insulin-like growth factor 1 (IGF-1) injections at days 4-6 following radiation results in restoration of saliva flow rates by day 30, making it a useful model to study mechanisms of repair(36). Mice were subjected to 5 Gy targeted head and neck IR and injected with IGF-1 via tail vein with either one dose at day 4 after IR for dissection and analysis at day 5, or three doses on days 4, 5, and 6 after IR and dissected for analysis by flow cytometry at day 30 post IR. Treatment with one dose of IGF-1 did not change total live immune cells at day 5 or day 30 when compared to IR-only controls (Fig 4A, Supplemental fig 6). Further, macrophages and neutrophils did not change at day 5 when given one IGF-1 injection compared to an IR only control; however, the increase in both macrophages and neutrophils seen at day 30 after IR was significantly attenuated with IGF-1 treatment (Fig 4B, Fig 4C Supplemental fig 6). We confirmed these data via immunofluorescent staining with an anti-F4/80 and anti-Ly6G antibody and analysis via manual counting (Fig 4D-G). These data indicate a restorative treatment attenuates the increase in neutrophils and macrophages seen at day 30 following IR, suggesting alterations in immune populations may play a role in the chronic loss of function phenotype. Control mice receiving IGF-1 treatment alone were injected with three doses of IGF-1 and parotid glands were collected 24 hours or 24 days after the last injection in order to mimic the day 30 IR time point. There were no significant changes in immune populations at day 24 after IGF-1 injections. We did observe a significant increase in CD4+ cells as a percentage of CD3+ T cells 24 hours after three injections with IGF-1. However, this was in part due to low cell number, with less than 20 cells contributing to the differences observed. Further, CD4+ T cells were not different from untreated at day 30 (Supplemental fig 7).

**Fig 4.**
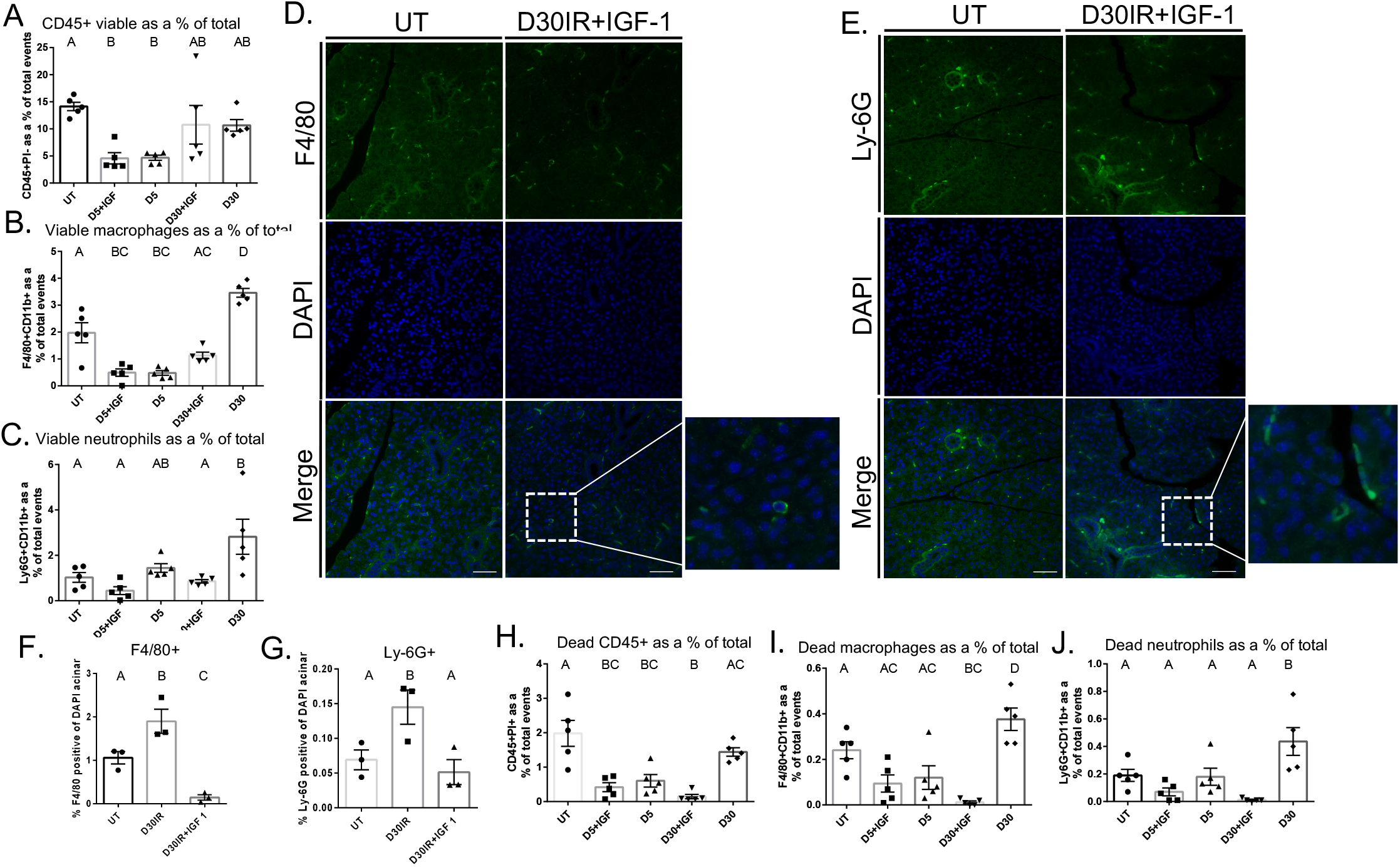
IGF-1 injections significantly attenuate the increase in macrophages and neutrophils observed at day 30 following IR. For analysis at day 5, IGF-1 was injected via tail vein at day 4 post IR, and parotid tissue was harvested 24 hours later. For analysis at day 30, IGF-1 was injected via tail vein at days 4, 5, and 6 following IR, and parotid tissue was dissected at day 30 post IR. Parotid tissue was prepared as a single cell suspension and labeled with antibodies for phenotyping via flow cytometry. A) Viable CD45+PI-total immune cells. B) Viable F4/80+CD11b+ macrophages. C) Viable Ly-6G+CD11b+ neutrophils. D) Immunofluorescent staining with anti-F4/80 antibody on FFPE salivary tissue. E) Immunofluorescent staining with anti-Ly6G antibody on FFPE salivary tissue. F) Quantification of F4/80 immunofluorescent staining determined by manual counting. G) Quantification of Ly-6G immunofluorescent staining determined by manual counting. H) CD45+PI+ immune cells. I) PI+ F4/80+CD11b+ macrophages. J) PI+ Ly6G+CD11b+ neutrophils. Scale bars represent 50 µM. Data analyzed by one-way ANOVA with Tukey’s post-hoc test and represented as mean ± SEM. Treatment groups with the same letter are not statistically different from each other.

**Supplemental Fig 6. Flow cytometry plots and gating strategy for phagocytic immune cells from IR damaged parotid glands in mice treated with IGF-1.** Flow cytometry data was analyzed manually via FlowLogic. Viable CD45+PI-gated from total events, F4/80+CD11b+ macrophages gated from CD45+PI-, Ly-6G+CD11b+ gated from CD45+PI-in: A-C) D5IR + IGF-1 mice, D-F) D30IR + IGF-1 mice.

**Supplemental Fig 7. IGF-1 injections alone do not alter the immune phenotype at day 30.** Mice were injected with IGF-1 via tail vein 3 times, 24 hours apart, and parotid tissue was dissected 24 hours after the last injection or at day 24 in order to mimic a day 30 time point. Flow cytometry data was analyzed manually via FlowLogic. Parotid tissue was prepared as a single cell suspension and labeled with antibodies for phenotyping via flow cytometry. A) Viable CD45+PI-total immune cells. B) Viable F4/80+CD11b+ macrophages, C) Viable Ly-6G+CD11b+ neutrophils. D) Viable CD3+ T cells. E) Viable CD3-cells. F) Viable CD49b+ NK cells. G) Viable CD4+ T cells. H) Viable CD8+ T cells. I) Viable CD4+CD8+ double positive T cells. J) Viable CD4-CD8-double negative T cells.

In mice treated with IGF-1, there was no significant difference in PI+ total immune cells at day 5 post IR, but there was a significant reduction PI+ total immune cells at day 30 versus IR only (Fig 4H). Moreover, treatment with IGF-1 did not change PI+ macrophages or PI+ neutrophils at day 5 when compared to IR-only controls; however, it significantly reduced PI+ macrophages and PI+ neutrophils at day 30 compared to untreated and an IR-only control (Fig 4I, 4J).

### IGF-1 injections attenuate the increase in CD4+CD8+ double positive T cells at day 30 post IR

Next, we aimed to characterize lymphocyte populations in the gland after IR and IGF-1 injections. IGF-1 injections did not change viable CD3+ T cells or CD3-cells at day 5 or day 30 compared to IR only controls (Fig 5A, 5B, Supplemental fig 5). IGF-1 administration did not change NK cells as a percentage of CD3-at day 5, but significantly reduced NK cells at day 30 versus an IR only control (Fig 5C, Supplemental fig 5). IGF-1 treatment did not change CD4+ T cells at any time point when compared to UT or IR controls (Fig 5D, Supplemental fig 5). Mice treated with IGF-1 injections had significantly increased CD8+ T cells as a proportion of CD3+ T cells at day 30 when compared to IR only mice (Fig 5E, Supplemental fig 5). Interestingly, IGF-1 injections did not change CD4+CD8+ T cells at day 5 but IGF-1 administration significantly attenuated the increase observed at day 30 in IR only mice (Fig 5F, Supplemental fig 5). IGF-1 administration did not change CD4-CD8-double negative cells at any time point versus untreated or IR only controls (Fig 5G, Supplemental fig 5).

**Fig 5.**
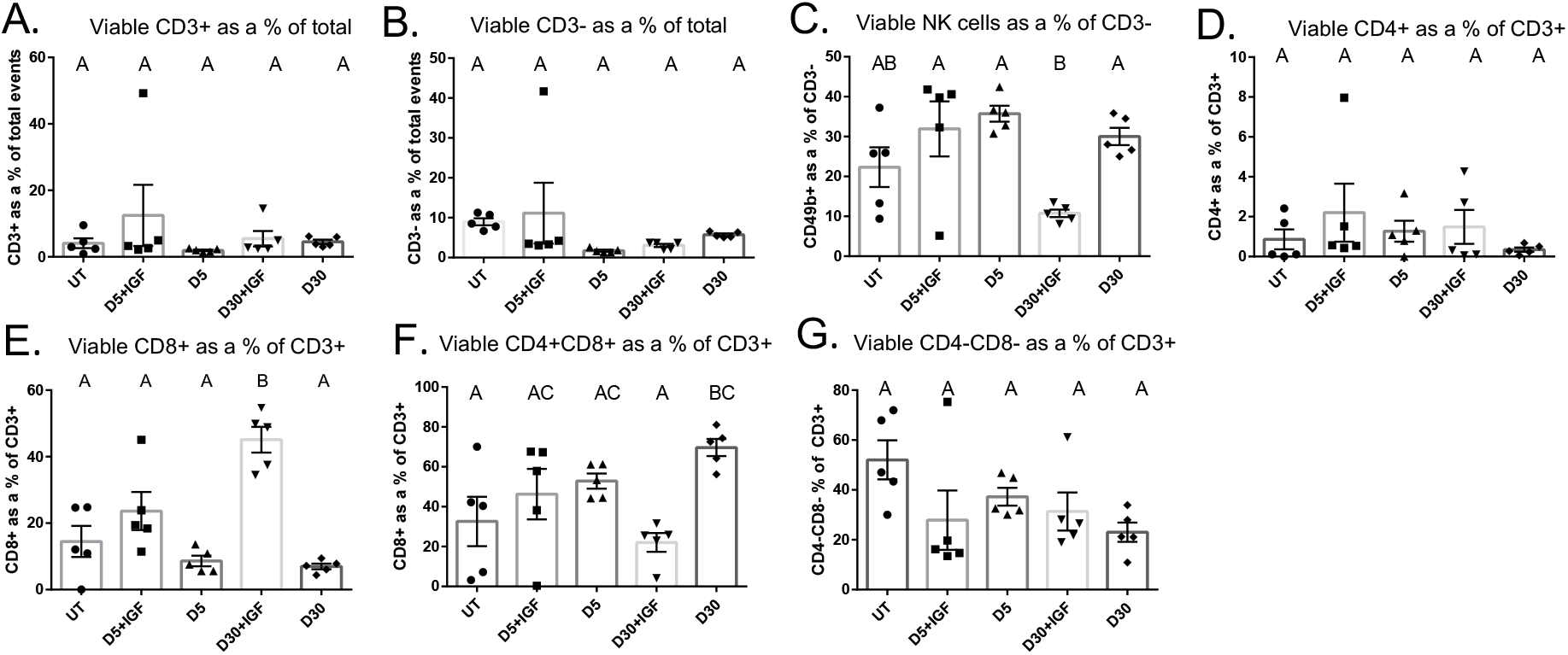
IGF-1 injections significantly attenuate the increase in CD4+CD8+ double positive T cells seen at day 30 after IR. For analysis at day 5, IGF-1 was injected via tail vein at day 4 post IR, and parotid tissue was harvested 24 hours later. For analysis at day 30, IGF-1 was injected via tail vein at days 4, 5, and 6 following IR, and parotid tissue was dissected at day 30 post IR. A) Viable CD3+ T cells. B) Viable CD3-cells C) Viable CD49b+ NK cells. D) Viable CD4+ T cells. E) Viable CD8+ T cells. F) Viable CD4+CD8+ double positive T cells. G) Viable CD4-CD8-double negative T cells. Data analyzed by one-way ANOVA with Tukey’s post-hoc test and represented as mean ± SEM. Treatment groups with the same letter are not statistically different from each other.

**Supplemental Fig 8. Flow cytometry plots and gating strategy for lymphocytes in IR damaged parotid glands from mice treated with IGF-1.** Flow cytometry data was analyzed manually via FlowLogic. CD3+ and CD3-gated from CD45+PI-, NK cells gated from CD3-cells, CD4+, CD8+, CD4-CD8-, CD4+CD8+ gated from CD3+ in: A-C) D5IR + IGF-1 mice, D-F) D30IR + IGF-1 mice.

### Pro- and anti-inflammatory cytokines decrease significantly at acute time points following radiation

Immune cells secrete cytokines as part of a normal inflammatory response and are increased in tissue following acute inflammation and injury(51, 52). Likewise, cytokines are typically increased in irradiated tissue(53). To better understand the inflammatory response in parotid glands following IR, we evaluated concentrations of cytokines in the salivary gland at days 2, 7, 14 and 30 post IR. Interestingly, all pro-inflammatory cytokines we analyzed, with the exception of IFN-γ, were significantly decreased at days 2 and 7 post-IR (Fig 6). Additionally, TNF-α, IL-1β, IL-23, IL-1ra, IL-27, IL-4, IL-17, and TIMP-1 were significantly decreased at day 14 when compared to controls but returned to untreated levels at day 30 (Fig 6A-H). Further, IL-2, IL-6, and IL-12 were significantly decreased at all time points following IR (Fig 6I-K). IFN-γ remained unchanged from untreated at days 2, 7, and 30, but was significantly decreased at day 14 (Fig 6L). Interestingly, IL-10, an anti-inflammatory cytokine, was significantly decreased at all time points evaluated (Fig 6M). These data show that cytokines are decreased acutely in parotid glands after IR, further substantiating that the immune and inflammatory response is dysregulated.

**Fig 6.**
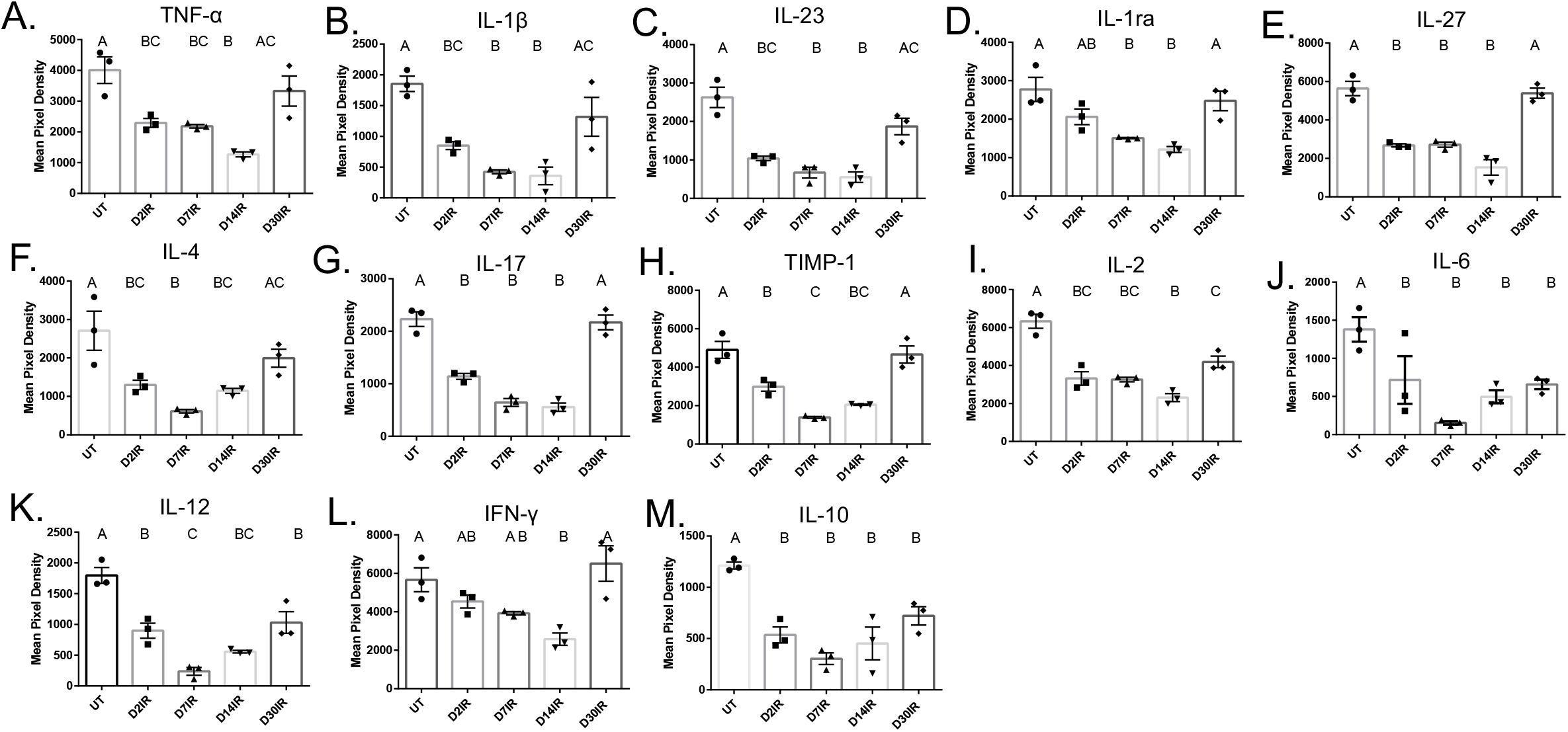
Pro- and anti-inflammatory cytokines are significantly decreased at acute time points following IR. Parotid tissue was dissected at days 2, 7, 14, and 30 following IR, and whole protein lysates were isolated and analyzed by cytokine array. A) TNF-α, B) IL-1β, C) IL-23, D) IL-1α, E) IL-27, F) IL-4, G) IL-17, H) TIMP-1, I) IL-2, J) IL-6, K) IL-12, L) IFN-γ, M) IL-10 analyzed by cytokine array. Data analyzed by one-way ANOVA with Tukey’s post-hoc test and represented as mean ± SEM. Treatment groups with the same letter are not statistically different from each other.

### Chemokine and chemoattractant concentrations significantly decrease in the parotid gland following IR treatment

Chemokines and chemoattractants are essential in leukocyte extravasation, and concentrations of chemokines increase significantly during wound healing(54, 55). Therefore, we quantified chemokines in parotid tissue to explore whether decreased chemokine concentrations could be resulting in insufficient immune cell infiltration. Concentrations of CXCL9 and CXCL11 were significantly decreased at days 2, 7, and 14 post IR, but returned to untreated levels at day 30 post IR (Fig 7A, B). CXCL10 was significantly decreased at days 7 and 14 but unchanged from controls at days 2 and 30 following IR (Fig 7C). CXCL2 was significantly decreased at all time points when compared to untreated (Fig 7D). CXCL1 appeared to decrease at day 14, but this was not significantly different from untreated (Fig 7E). Finally, we measured levels of macrophage chemoattractants in the gland following IR. Remarkably, GM-CSF was significantly decreased at all time points (Fig 7F). M-CSF significantly decreased at day 14 but returned to untreated levels at day 30 (Fig 7G). Interestingly, MCP-1 did not change at any time point following IR (Fig 7H). Finally, MCP-5 significantly decreased at days 2, 7, and 14 following IR, and recovered to untreated levels at day 30 (Fig 7I). These data suggest an insufficient chemokine response which may be contributing to the abnormal immune infiltrate observed, and this could further indicate dysregulation of the immune response in the parotid glands after IR and may be contributing to dysfunction.

**Fig 7.**
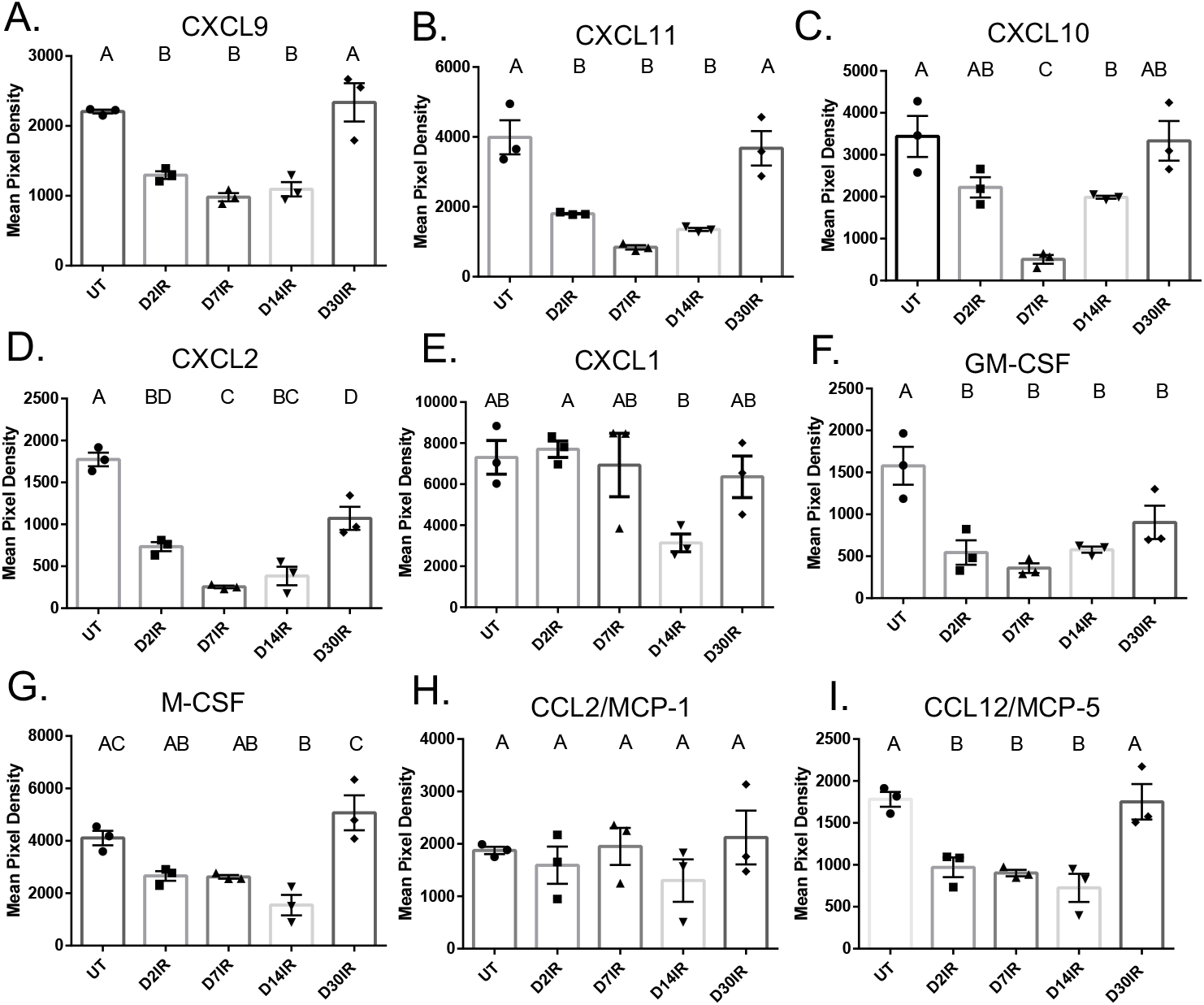
Chemokines, with the exception of CXCL1 and MCP-1, decrease acutely following 5 Gy IR. Parotid tissue was dissected at days 2, 7, 14, and 30 following IR, and whole protein lysates were isolated and analyzed by cytokine array. A) CXCL9, B) CXCL11, C) CXCL10, D) CXCL2, E) CXCL1, F) GM-CSF, G) M-CSF, H) CCL2/MCP-1, I) CCL12/MCP-5 analyzed by cytokine array. Data analyzed by one-way ANOVA with Tukey’s post-hoc test and represented as mean ± SEM. Treatment groups with the same letter are not statistically different from each other.

## Discussion

Salivary gland dysfunction after radiation therapy is a serious side effect for head and neck cancer patients that often persists years after treatment and results in diminished quality of life (56). Currently, long-term treatments are not commonly prescribed due to adverse side effects, short-term treatments are often ineffective and only target the symptoms, and there is no standard of care for restoration of function. In the present study, we aimed to characterize the immune and inflammatory response in parotid salivary glands after radiation. We quantified and phenotyped select immune cell populations and measured cytokine concentrations in the parotid glands following targeted head and neck radiation. We determined that total immune cells and macrophages are decreased in the parotid glands at days 2 and 5, and that neutrophils, macrophages, and CD4+CD8+ T cells are significantly increased at day 30 (Fig 1, Fig 3). In addition, inflammatory cytokines decrease significantly at acute time points (Fig 6, Fig 7). Further, we found that injections with IGF-1, which restore salivary gland function at day 30, attenuate the increases observed in macrophages, neutrophils, and CD4+CD8+ T cells at day 30 (Fig 4, Fig 5). These data suggest irradiated parotid glands do not have sufficient acute phagocytic cell infiltration and cytokine production that is normally observed following tissue injury, which may contribute to prolonged glandular dysfunction after radiation.

Wound healing is a complex, active response to injury and comprises inflammatory, proliferative, and remodeling phases, and immune cells are involved in each of these three phases. In models of wounds that heal normally, there is significant neutrophil and macrophage infiltration at acute time points(57). Additionally, radiotherapy can dampen the immune response through apoptosis of immune cells(58); however, neutrophil and macrophage infiltration into tissue after radiation is observed in multiple tissue types(9, 10, 42). Our data indicate that macrophages and total immune cells are decreased acutely in parotid tissue following a single dose of 5 Gy IR. Several studies have suggested that CD45 expression increases in tumors in vivo after radiation treatment(59–61). However, it is well documented that CD45 expression is directly correlated to the number of immune cells in the tumor and thus, the increase observed is due to increased immune infiltration in the tumor(62). A recent study by Zhao et al. found that 15 Gy IR significantly downregulated mRNA expression of phagocytosis receptors Fcgr1 and Axl and reduced macrophage populations in the submandibular gland, corroborating our findings(44). This study also reported that Shh gene transfer restored salivary gland function while restoring macrophage populations, suggesting an important role for macrophages in healing. Further, the same group found that activation of Hedgehog signaling restored glandular function and preserved mRNA markers of macrophages in parotid glands, thereby inhibiting cellular senescence and inflammation, further indicating that macrophages are necessary in glandular regeneration(50). Phagocytosis is vital to tissue healing and restoration of function, thus, the decrease in phagocytosis and phagocytic cells may be contributing to loss of function and the inability of salivary glands to regenerate. In other tissues that undergo improved healing following IR, including the intestine and lung, there is a significant increase in macrophages following IR, thus suggesting macrophage infiltration is a vital response to IR injury (9–11). Apoptosis of neutrophils signals the tissue to suppress the inflammatory phase by downregulating the expression of pro-inflammatory cytokines and increasing expression of anti-inflammatory cytokines, thereby facilitating the transition between the inflammatory and the proliferative phase of wound healing(63, 64). Neutrophils in the parotid gland appear to infiltrate later than anticipated, and do not seem to be undergoing apoptosis as expected; we expected to see an increase in PI+ neutrophils at day 2 after IR due to apoptosis of acinar cells occurring 24-48 hours after IR(65). Proliferation is an important component of the wound healing response; however, compensatory proliferation of the acinar compartment of parotid glands is a persistent phenotype that begins at day 5 after IR and continues through day 90(35). Several studies have reported that the elevated proliferative response in salivary glands is highly correlated with loss of function and treatments that restore salivary flow also attenuate compensatory proliferation of acinar cells(35, 37, 66). Due to the fact that we did not observe an increase in neutrophil apoptosis, we postulate the parotid gland does not receive the necessary signals to transition from the inflammatory phase to the proliferative and regenerative phases, therefore contributing to the abnormal proliferative response and the lack of regeneration and restoration of function. Because we observed significant decreases in total viable immune cells and macrophages at days 2 and 5, we hypothesize treatments that restore populations of phagocytic immune cells at acute time points may improve healing by attenuating the proliferative response. The impact of the immune system on salivary gland function as well as the underlying mechanisms warrant further investigation.

In addition to phagocytic immune cells, lymphocytes play an important role in healing and tissue regeneration; they are involved in regulating the cellular response to injury and are thought to control the balance between a regenerative or profibrotic outcome(25). Lymphocytes are recruited to the site of injury by chemokines and are the last immune cells to infiltrate the wound; additionally, some studies suggest they accumulate in radiation damaged tissue at both acute and chronic time points(29). It is well documented that lymphocytes are highly radiosensitive and radiotherapy consistently causes severe and persistent peripheral lymphopenia; however, our understanding of the role of lymphocytes in the radiation damage response in tissue is limited(67, 68). We expected to see significant decreases in total viable lymphocytes and increases in PI+ lymphocytes following IR. Interestingly, only CD3-cells were decreased at acute time points, and there were no significant increases in PI+ lymphocytes of any population at any time point measured. CD4+ T cells are the most abundant lymphocytic cell type in wound healing, and surprisingly, this cell type represented only ∼1% of all CD3+ at all time points with little change after radiation. Unexpectedly, there was a significant increase in double positive CD4+CD8+ T (DPT) cells at day 30 after IR. DPT cells are most often described in pathological conditions, and the role of DPT cells is largely understudied and not well understood. The literature has shown both cytotoxic and immunosuppressive functions for DPT cells in cancers, autoimmune diseases, and HIV(69–72). The increase in DPT cells in parotid glands at day 30 is not observed in the IGF-1 restorative model, so this may imply a compensatory response to the lack of regeneration. Interestingly, a study utilizing single cell RNA sequencing (scRNAseq) found that fractionated IR resulted in a significant increase in DPT cells at 10 months following treatment, and further, reported that DPT cells had the highest number of radiation-induced differentially expressed genes(73). This paper also reported substantial immune-epithelial cell interactions, specifically increased signaling from acinar to DPT and CD8+ T cells. Additional studies are necessary to ascertain the role of DPT cells in parotid glands following IR.

An inflammatory response following an injury is thought to be a prerequisite to complete wound healing. Previous data indicate that treatment with indomethacin, a COX inhibitor, prior to IR does not improve salivary flow rates, but treatment at days 3, 5, and 7 after IR restores salivary flow rates at day 30(33). Our data show that concentrations of both pro and anti-inflammatory cytokines are significantly decreased in the parotid gland following IR, further suggesting salivary glands are undergoing a dysregulated inflammatory response after radiation, as a significant increase in pro-inflammatory cytokines is observed in multiple tissues following IR(8, 53, 74–77). Certain pro-inflammatory mediators, such as IL-6, TNF-α, and IL-1β are vital to proper wound healing in an appropriate window. Multiple studies show that mice lacking IL-6 exhibit deficiencies in wound healing, and we postulate the decrease in pro-inflammatory cytokines following radiation, including IL-6, may contribute to the lack of regeneration(22, 23). Moreover, an increase in anti-inflammatory cytokines, such as IL-10, is typically observed at the end of the inflammatory phase in order to help restore tissue homeostasis and progress wound healing towards redifferentiation. Absence of IL-10 can result in increased monocyte infiltration and expression of pro-inflammatory cytokines(24). We speculate that the decrease observed in IL-10 in parotid glands at all time points following IR is contributing to lack of regeneration; however, the data are contradictory and has shown that IL-10 can have inhibitory or contributory mechanisms on wound healing and fibrosis(78–80). The significant decreases observed in IL-10 through day 30 post IR may indicate a chronic inflammatory response at later time points, which is corroborated by increased T cells and robust differential gene expression in these cells reported in single cell RNAseq data. Additionally, our data show cytokine concentrations and immune cells, particularly macrophages, follow the same trend after radiation; however, it is unclear if immune cells are migrating out of the gland following IR leading to lower cytokine production, or if decreased cytokine and chemokine concentrations are resulting in fewer immune cells being attracted to the gland. We hypothesize that IR-damaged acinar epithelial cells are unable to mount a substantial chemokine and cytokine response in order to stimulate an appropriate immune response. Future studies are warranted to gain understanding on the role of cytokines and chemokines in the parotid gland following radiation.

Previous studies have reported that injections with IGF-1 after IR restore salivary flow rates through attenuation of compensatory proliferation and activation of aPKCζ, and the present study showed that IGF-1 injections alter the immune phenotype at day 30 following radiation(36). Interestingly, IGF-1 did not change total immune cells, macrophages, neutrophils, or lymphocytes at day 5 following IR. Treatment with IGF-1 maintains activation of aPKCζ and attenuates compensatory proliferation in the salivary gland, which are correlated with restoration of function(35, 36), and administration of IGF-1 attenuated the increase seen in macrophages neutrophils, and DPT cells at day 30. Expectedly, IGF-1 reduced PI+ total immune cells, macrophages, and neutrophils in the parotid gland at 30 days post IR. Currently available data, though limited, suggests immune cells may be crucial for restoration of salivary gland function, and our results substantiate these data (44, 50). We postulate that the acute immune response influences the chronic response and is underlying prolonged loss of function, and administration of IGF-1 results in parotid tissue undergoing a more typical healing timeline as defined by what is observed in the literature, resulting in reductions in proliferation and the altered immune phenotype seen at day 30. It is unclear if IGF-1 itself is driving the immune alterations we observed at day 30, or if the attenuation of proliferation and activation of aPKCζ induced by IGF-1 could be indirectly impacting the immune response. Our data does not elucidate the mechanism by which IGF-1 is altering the immune response; therefore, future work should utilize the IGF-1 model in cytokine or macrophage deficient mice as well as explore the mechanism of the direct effects of IGF-1 on immune cells.

In conclusion, these data suggest the parotid gland does not progress through a normal inflammatory response and does not have the expected immune phenotype after radiation damage, and this is likely contributing to insufficient wound healing and prolonged loss of function. Future directions include investigating the mechanisms contributing to the abnormal immune phenotype in salivary glands and utilizing immune deficient mice to observe if certain populations are required for restoration of function. Importantly, understanding how the inflammatory response and immune phenotype impact lack of regeneration in the parotid gland could help uncover new potential therapeutic treatments to permanently restore salivary gland function.

## Acknowledgements

We would like to acknowledge Dr. Douglass Diak, Dr. Forrest Baker, and Dr. Kyle Smith for their experimental help with flow cytometry and data analysis. We would like to acknowledge Dr. Brenna Rheinheimer for assistance in immunofluorescence staining. We would like to acknowledge Dr. Sean Limesand for the use of the Leica microscope.

## Notes

### Competing Interest Statement

The authors have declared no competing interest.

